# Thermal Plasticity of Stage-specific Development Time and Adult Body Size under Temperature Shifts: A Case Study Using *Drosophila melanogaster*

**DOI:** 10.1101/2024.11.16.623922

**Authors:** Aradhya Chattopadhyay, Rishav Roy, Payel Biswas, Shampa M. Ghosh

**Affiliations:** School of Biotechnology, Kalinga Institute of Industrial Technology (KIIT), Bhubaneswar, India; School of Biological Sciences, University of Edinburgh, UK

**Keywords:** thermal plasticity, life stage-specific thermal plasticity, development time, thermal sensitivity, thermal shift, temperature-induced developmental plasticity, body size, *Drosophila*

## Abstract

We examined how thermal shifts influence development time and adult body size in *Drosophila melanogaster*. Individual flies were exposed to alternating temperatures of 25°C (optimal) and 17°C (cold), with shifts introduced at key developmental transitions: larval hatching and pupariation. We found while larval-stage temperature is the biggest determinant of thermal plasticity of development time and adult size, the egg-stage temperature also influences the pace of development and growth throughout pre-adult duration. The effect of low-to-high and high-to-low temperature shifts on development and growth may not be symmetric. When eggs are reared at 25°C and then shifted to 17°C, larval and pupal durations undergo reduction compared to constant 17°C, but it produces slightly larger adults. A higher egg-stage temperature thus seem to exert a carryover effect that accelerates subsequent development and growth when later stages experience colder temperatures. Surprisingly, flies whose egg stage is exposed to 17°C followed by a shift to 25°C also have reduced larval duration and larger size, relative to those developing at constant 25°C. We speculate this could be either because 17°C to 25°C represents a low-to-high temperature shift or a sub-optimal-to-optimal thermal shift that results in metabolic and/or hormonal changes accelerating differentiation and growth. While pupal duration is sensitive to current and to some extent prior thermal environments, it does not contribute substantially to thermal plasticity of size. Development time is longer in males than in females, and this difference seems to start from larval stage while the pupal duration plays a bigger role in creating this sex-specific difference. Overall, employing individual fly rearing, this study helped to unravel the effect of thermal shifts on growth and development in *D. melanogaster* with great precision.

## Introduction

Understanding how climate change affects organisms and shapes their adaptive responses are critical questions in thermal biology. Temperature influences a wide array of biological processes, including development, physiology, metabolism, behaviour, and life history (Huey and Kingsolver, 1989; Johnston and Bennett, 1996; Angilletta, 2009). While natural environments inherently exhibit temperature variability, organisms vary in their susceptibility to thermal changes (Angilletta, 2009). Ectotherms, in particular, are highly vulnerable due to their reliance on ambient temperatures to regulate body function (Addo-Bediako et al., 2000; Angilletta, 2009). Consequently, changes in environmental temperature can have immediate and profound effects on their performance and fitness (Huey and Stevenson, 1979; Pinsky et al., 2019). These effects are manifested through thermal plasticity of biological traits. In recent years, there has been growing interest in understanding mechanistic regulation and adaptive significance of thermal plasticity, as it reflects organismal responses to climate change (Trotta et al., 2006; Cooper et al., 2012; McDonald et al., 2018; Lafuente et al., 2024).

While research on thermal plasticity has traditionally focused on examining how various traits respond to a range of constant temperatures under controlled laboratory conditions (Trotta et al., 2006; Klepsatel et al., 2019; Lafuente et al., 2024), attention has also been paid to understanding how organisms respond to thermal fluctuations, which are intrinsic features of natural environments. Studies have investigated the effects of such fluctuations on development, performance, and life history traits (Al-Saffar 1995, 1996; Petavy et al., 2001; Manenti et al., 2015, 2021; Tobler et al., 2015) However, in addition to daily and seasonal variation, temperature can also differ across life stages, particularly in species with complex life cycles. For example, holometabolous insects go through different life forms during their development, each of which may occupy different microhabitats and thus may experience varying temperature and humidity (Taboada et al., 2013; Woods, 2013; Rolff et al., 2019; Kingsolver and Buckley, 2020; Vives-Ingla et al., 2023). Despite this, we know relatively little about how such stage-specific thermal variability influences developmental plasticity across the ontogeny.

*Drosophila*, a widely distributed ectothermic holometabolous insect, serves as an excellent model for investigating thermal plasticity. This is due to its broad latitudinal range, which reflects significant natural variation in temperature exposure (James et al., 1997; Trotta et al., 2006; Castañeda et al., 2015), and its experimental tractability, particularly the ease with which temperature conditions can be precisely manipulated in the laboratory (Huey et al., 1991; Partridge and French, 1996; Ghosh et al., 2013; Kellermann and van Heerwaarden, 2019). Life cycle of *Drosophila* consists of four distinct stages: egg, larva, pupa, and adult. In natural environments, these stages may occupy different microhabitats and, consequently, diverse thermal conditions (Lockwood et al., 2018; Moghadam et al., 2019). The egg stage is immobile and typically laid on the surface of fermenting or rotting fruit. The larval stage is motile but largely confined within the fruit substrate, where temperature and humidity can be buffered (Evans et al., 2017; Schöneberg et al., 2020). As larvae mature, they exit the fruit to search for a suitable pupariation site, often exposing themselves to more variable environmental conditions (Sokolowski et al., 1986; Powell, 1997). In contrast, adults are fully mobile and can actively thermoregulate through behaviour (Reviewed in Dillon et al., 2012). These differences in spatial and behavioural ecology suggest that the thermal environment experienced by each ontogenetic stage can vary substantially due to factors such as substrate differences, daily temperature cycles, or extreme thermal events like heatwaves (Kingsolver et al., 2011; Moghadam et al., 2019; Schöneberg et al., 2020).

Previous research has shown that thermal tolerance and reaction norms can differ across life stages, implying that certain stages may be more thermally sensitive than others (e.g. Kingsolver et al., 2011; Austin and Moehring, 2013; Blanckenhorn et al., 2014; van Heerwaarden et al., 2014;). Building on these findings, we hypothesize that exposure to thermal variation during development will have stage-specific effects, potentially altering the duration of individual pre-adult stages and influencing final adult body size. Understanding these differential effects is critical for uncovering the mechanisms by which thermal environments shape developmental trajectories and life history outcomes in ectothermic organisms.

Thermal plasticity of developmental time has been extensively studied in *Drosophila*, and like most ectotherms, flies reared at lower temperatures exhibit prolonged development (Davidson and Davidson, 1944; Partridge et al., 1994b; Al-Saffar et al., 1995; Ghosh et al., 2013). Developmental events during embryogenesis scale uniformly across temperature, such that the sequence and proportional timing of events remain consistent even as the total duration changes (Kuntz and Eisen, 2014; Chong et al., 2018). Similarly, pre-adult stages including larval instars and the pupal period exhibit a proportionate increase in duration as temperature decreases (Ghosh et al., 2013). The optimal rearing temperature for *Drosophila melanogaster* is approximately 25°C, whereas development at 17°C takes nearly twice as long. All key developmental stages, that is, egg hatching, larval growth, and pupation, are extended roughly two-fold at 17°C compared to 25°C (Ghosh et al., 2013).

While much is known about developmental plasticity under constant warm or cold temperatures (Huey et al., 1995; Ghosh et al., 2013; Fallis et al., 2014; Klepsatel et al., 2019; Szabla et al., 2024), and under daily temperature fluctuations (Le Vinh Thuy et al., 2016; Saxon et al., 2018; Manenti et al., 2021, 2015), relatively little is understood about how *Drosophila* development responds to thermal shifts occurring across developmental transitions. For example, what happens if temperature changes after larvae hatch from the eggs, or between larval and pupal stages? Does the timing of the shift matter? Does development continue to scale proportionally, or are certain stages more sensitive to changes than others? In light of these questions, we sought to investigate how stage-specific temperature shifts influence the subsequent development of the fly. In this study, we investigated how temperature shifts across different developmental stages, specifically egg, larva, and pupa affect the duration of subsequent stages and the overall development time of *Drosophila melanogaster*. Our goal was to identify the presence and nature of carryover effects, wherein thermal conditions experienced during an earlier developmental stage influence the duration of later stages. We asked: If a developmental stage is reared under relatively cold conditions but preceded by a warmer stage, does the prior exposure to warmer temperature accelerate the current stage’s development? Conversely, does exposure to colder temperature during an earlier stage impose a developmental lag, even when the current stage is reared under optimal warm conditions? Additionally, we examined whether these carryover effects vary across the sexes.

Like development time, ectothermic growth is heavily reliant upon ambient temperature (Zuo et al., 2011; Ghosh et al., 2013; Kingsolver et al., 2015; Garrad et al., 2016). Pre-adult development and growth are major determinants of insect body size, as the adults have a hard exoskeleton that limits their capacity to grow (Simpson et al., 1980; Nijhout et al., 2006; Shingleton, 2010; Hanna et al., 2023). While factors like nutrition, age, and reproductive investment may lead to weight fluctuations in an adult insect to some extent (Gerofotis et al., 2019; Poças et al., 2022), size is principally contingent upon the growth achieved during its pre-adult development. 85% ectotherms grow bigger when they develop in colder environments. This inverse relationship between temperature and body size is known as the temperature size rule (TSR) (Angilletta et al., 2004; Walters and Hassall, 2006; Einum et al., 2021). Temperature size rule is well-documented in *Drosophila* (David et al., 1994; French et al., 1998; Ghosh et al., 2013; McDonald et al., 2018). The possible fitness consequence of TSR and its developmental basis have been topics of intense debate for decades (Atkinson, 1994; Van Der Have and De Jong, 1996; Kingsolver and Huey, 2008). Larger body size in cooler environments may be advantageous because lower surface area-to-volume ratios reduce relative heat loss and enhance thermal retention (Bergmann 1847, Mitchell et al., 2018). However, the fitness consequences of TSR might be more complex than that, and a clear understanding of this phenomenon is still lacking. The developmental basis of TSR has been explored to some extent through empirical studies (Van Der Have and De Jong, 1996; Gibert and De Jong, 2001; Angilletta and Dunham, 2003; Ghosh et al., 2013). Higher temperature generally leads to an increase in both developmental and growth rates (French et al., 1998; Gillooly et al., 2001). Consequently, warmer temperature may make an organism grow faster in size but it would also reduce the duration of growth as the organism completes its development earlier than in colder temperatures (Zuo et al., 2011; Ghosh et al., 2013; Semsar-kazerouni et al., 2022). This is thought to be the underlying reason for ectotherms developing a smaller body size in warmer temperatures (Van Der Have and De Jong, 1996; Atkinson and Sibly, 1997; Angilletta et al., 2004; Kingsolver et al., 2007). However, the mechanistic basis of TSR may be more nuanced than this and it may vary across species and thermal ranges (Davidowitz and Nijhout, 2004; Ghosh et al., 2013).

Thermal regulation of size in *Drosophila* is often studied either at a range of constant developmental temperatures (Partridge et al., 1994a; Nunney and Cheung, 1997; Ghosh et al., 2013; Lafuente et al., 2018; Klepsatel et al., 2019), or environments with daily thermal fluctuations (Kjærsgaard et al 2012; Manenti et al., 2015, 2021; Saxon et al., 2018). In contrast, how growth and plasticity are regulated when temperature varies across different developmental stages remains largely unexplored. The reason for the same is flies are typically grown in groups cultures, making it difficult to track the effects of changing temperatures on development in a stage-specific manner, because not all flies undergo transition synchronously between stages. In a study by Hoover and Marks, 2021, eggs were reared at 18°C and 28°C and cultures were shifted to alternate temperatures after 24 hours. The study showed that adult size in *Drosophila* is contingent upon early developmental temperature. Nonetheless, a comprehensive understanding of how thermal changes across developmental stages shape body size plasticity, and whether these effects differ between male and female flies, remains unclear.

In view of the above, we devised a unique way and investigated how the temperature experienced during one stage influences growth during the subsequent stages and how variations in the thermal environment across different pre-adult developmental stages affect the final body size of flies. We employed a novel experimental design in which individual *Drosophila melanogaster* eggs were placed in microcentrifuge tubes containing food shortly after oviposition, then subjected to temperature shifts at key developmental transitions-either at hatching (egg-to-larva), pupariation (larva-to-pupa), or both. We used two contrasting but biologically relevant rearing temperatures: 25°C (optimal) and 17°C (cold), enabling us to implement both low-to-high and high-to-low temperature shifts. By introducing temperature changes at specific developmental transitions, we aimed to disentangle stage-specific versus cumulative effects of thermal conditions on developmental trajectories. This setup allowed precise measurement of stage-specific durations of males and females under each temperature condition and the body weight of individual flies upon emergence.

## Materials and Methods

### Population ancestry

We used a large, outbred laboratory population of *Drosophila melanogaster* named CO-mix for this study. It is a descendant of the ancestral CO-populations (Rose et al., 1984; Chippindale et al., 1993, 1997). 5 replicates of CO populations were mixed to create the CO-mix population. CO populations and their ancestors are being maintained at 25°C for decades. CO-mix flies are maintained at a population size of above 1200 flies, on a 14-day discrete generation cycle, at 25°C under 24 hours light, and ∼70% relative humidity on standard cornmeal medium. Rearing density is controlled at ∼70 eggs per 6 ml food in a vial. Twenty-five such vials are reared every generation for growing the flies, and upon eclosion, all flies are transferred to a population cage, and the eggs are collected after 3 days and incubated in vials, to initiate the next generation. At the time of the assay discussed in the paper, CO-mix populations were 15 generations old.

### Experimental protocol

For assaying stage specific development time and body size of males and females, individual flies were grown in microcentrifuge tubes (MCTs), each containing 300 µL of cornmeal food. *Drosophila* development includes three major developmental transitions, namely egg to larva (hatching), larva to pupa (pupariation), and pupa to adult (eclosion). For different thermal treatments, thermal shifts between the two treatment temperatures, namely, 17°C and 25°C, were performed either at hatching, or at pupariation, or at both. Total eight thermal combinations were used for the experiment, as shown in Table 1.

**Table 1:** Schematic of the 8 thermal treatments.

| Treatment | Egg | Larva | Pupa | Sample size (N) |
| --- | --- | --- | --- | --- |
| 1 | 25°C | 25°C | 25°C | 48 |
| 2 | 25°C | 17°C | 25°C | 43 |
| 3 | 25°C | 17°C | 17°C | 53 |
| 4 | 25°C | 25°C | 17°C | 47 |
| 5 | 17°C | 17°C | 17°C | 68 |
| 6 | 17°C | 17°C | 25°C | 69 |
| 7 | 17°C | 25°C | 17°C | 70 |
| 8 | 17°C | 25°C | 25°C | 71 |

A small agar piece containing a single egg was placed in each MCT within 2 hours of egg lay from the CO-mix populations. In *D. melanogaster*, eggs appear smooth and swollen when larvae are inside, and after larval hatching, the eggs appear crumpled. This way hatched vs. unhatched eggs can be distinguished clearly under a microscope. For treatments in which eggs were reared at 25°C, eggs were monitored for larval hatching 12 hours onwards post egg collection, at every 2-hour intervals. The time of larval hatching was noted down for each MCT. Hatching check was continued till 26 hours post egg collection, when no new eggs hatched for 3 consecutive checks. Eggs that failed to hatch and became blackened, were discarded. For treatments that required a downshift, individual tubes were shifted from 25°C upon hatching and incubated at 17°C. For the remaining, MCTs were maintained at 25°C. Similar protocol was followed for eggs reared at 17°C, and larval hatching was monitored from 32 hours up to 42 hours post egg collection. For treatments requiring upshifts, MCTs were shifted from 17°C to 25°C upon larval hatching, while for the other treatments, MCTs were maintained at 17°C during the larval stages.

Hatched larvae were allowed to grow at specific temperatures and subsequently monitored for pupariation after few days. MCTs were monitored for formation of pupae on inner wall of the tube or lower side of the tube’s lid every 4 hours. Upon pupariation, the time was noted, and tubes were retained at 25°C or 17°C, for treatments that involved no shifts at pupariation. Checks were continued till no new pupa was observed for 3 successive checks. Larvae that failed to reach the wandering stage or pupariate, were discarded. For treatments involving shifts at pupariation MCTs with pupae were shifted to specified temperature after noting their pupariation time. As metamorphosis ensued over next few days and pupae became dark indicating development of adult flies within pupal cases, individual tubes were monitored for eclosion. As eclosion started, checks were continued every 4 hours and time was noted along with sex of the flies. Upon eclosion, flies were freeze-killed by keeping them at -20°C for 40-45 mins and subsequently weighed using a Sartorius Quintix 35 (*d* = 0.01mg) balance. Eclosion checks were continued till no new flies ecolsed for 3 successive checks.

For each thermal treatment, number of MCTs used ranged from 60 to 80, each containing 300 µl of cornmeal food. Initially we had anticipated survivorship of flies will be lower at 17°C compared to 25°C as cold temperature has been shown to reduce survivorship in several studies (McKenzie, 1975; Amarasekare and Sifuentes, 2012; Klepsatel et al., 2019). Therefore, we used 60 and 80 eggs in treatments starting with 25°C and 17°C respectively. We observed survivorship above 80% in all four treatments initiated at 17°C and in two of the treatments initiated at 25°C. In the remaining two 25°C treatments, survivorship was lower (72% and 78%), likely due to stochastic variation. The high survivorship in the 17°C treatments may reflect some unknown aspect of fly rearing in MCTs. As a result, sample sizes ranged from 43 to 71 across treatments.

### Data analysis

For a given MCT, egg duration was calculated by subtracting the midpoint of the 2 hour-long egg collection window from time of larval hatching. Larval duration for each fly was calculated by subtracting hatching time from the time of pupariation. Pupal duration was calculated by subtracting pupariation time from the time of eclosion. Total egg to adult development time was also noted for individual flies in all 8 thermal treatments. Wet weight of individual flies was noted.

All data, except those excluded based on predefined criteria (for example, failed eclosion or incomplete developmental records), were included in the analysis. Prior to statistical testing, each dataset was evaluated for normality using Shapiro-Wilk tests (Shapiro and Wilk, 1965) and visual inspection of Q-Q plots (Supplementary figures S1-S5). Because development time and wet weight measurements consistently violated assumptions of normality and homoscedasticity across multiple treatment groups, all analyses were conducted using nonparametric methods. Traditional nonparametric tests such as Kruskal-Wallis or Friedman test are not suitable for the experimental design used in this study as both tests are limited to single-factor comparisons and cannot model multi-way interactions between independent variables. Because our dataset includes a four-factor full factorial structure (egg, larval, and pupal temperatures, and sex), along with biologically meaningful interactions among these stages, an analysis framework capable of testing interaction terms was required. Therefore, we employed a full factorial aligned rank transform (ART) ANOVA for the analyses using r-package ARTool (Wobbrock et al., 2011; Elkin et al., 2021; Kay et al., 2025). ART was chosen because it enables nonparametric testing of multi-factorial designs while preserving interpretability of interaction effects. For egg duration effect of egg temperature, sex and their interaction were tested. For larval duration effect of egg and larval temperatures, sex, and interaction of the three factors were tested. For the remaining three variables, namely, pupal duration, egg to adult development time, and wet weight of flies, four-factorial ART ANOVAs were performed with egg, larval and pupal temperatures and sex as factors.

Post hoc pairwise comparisons were performed using estimated marginal means (emmeans) (Lenth and Piaskowski, 2017) computed on ART-transformed data, with p-values adjusted for multiple comparisons. All tests were two-sided unless indicated otherwise. No transformations were applied to the response variables prior to nonparametric testing. ART-ANOVA results including degrees of freedom, sum of squares, F-values, and adjusted p-values for factorial and pairwise analyses are reported in the Results section. Data analysis was performed using custom R scripts written for this study.

## Results

### Development Time

At 25°C, mean egg duration (hatching time) of flies is 21 h, while at 17°C eggs take 36 h to hatch on average (Supplementary figure S6), reflecting a 1.7-fold change in egg duration. Mean larval development time changes from 95 h to 196 h, and mean pupal development time increases from 96 to 201 h when flies are grown at 17°C as opposed to 25°C. Thus, larval and pupal durations extend by two-fold from 25°C to 17°C.

### Egg and larval durations

17°C makes egg and larval durations significantly longer compared to 25°C (*p* < 0.0001). Egg duration does not differ across the sexes (Table 2).

**Table 2:** ANOVA table from aligned rank transform (ART) analysis showing nonparametric factorial effects of egg temperature and sex on egg duration.

| *Effect* | *df* | *df.res* | *SS* | *F* | *P* |
| --- | --- | --- | --- | --- | --- |
| Egg temperature | 1 | 465 | 6213590 | 1543.12 | <0.0001 |
| Sex | 1 | 465 | 14851.5 | 0.86047 | 0.35409 |
| Egg temperature × Sex | 1 | 465 | 4395.58 | 0.24968 | 0.61753 |

Under constant temperature rearing, average larval duration at 25°C is 95 h, and at 17°C, it is 196 h. Egg and larval-stage temperatures show significant interaction in shaping larval duration (*p* < 0.0001) (Figures 1a, 1b) (Table 3). Downshifting temperature from 25°Cat egg-stage to 17°C larval temperature (25°Cegg-17°Clarval) significantly reduces mean larval duration (181 h) relative to larval development time observed at constant 17°C (196 h) (*p* < 0.0001). Interestingly, upshifting temperature from 17°C to 25°C from egg to larval stage (17°Cegg-25°Clarval) also causes a small but significant reduction in mean larval duration (90 h) compared to larvae experiencing 25°C during both egg and larval stages (95 h) (*p* < 0.0001). ART ANOVA results indicate significant effect of sex (*p* < 0.0001), egg temperature × sex (*p* < 0.0001), and larval temperature × sex (*p* = 0.0268) interactions. The average difference between male and female larval duration pooled over different treatments is 0.9 h. Males have a slightly longer larval duration (0.5 to 4 h slower) than females across different thermal treatments.

**Table 3:** ANOVA table from aligned rank transform (ART) analysis showing nonparametric factorial effects of egg temperature, larval temperature and sex on larval duration.

| *Effect* | *df* | *df.res* | *SS* | *F* | *P* |
| --- | --- | --- | --- | --- | --- |
| Egg temperature | 1 | 461 | 620849 | 38.4675 | <0.0001 |
| Larval temperature | 1 | 461 | 6114328 | 1323.02 | <0.0001 |
| Sex | 1 | 461 | 463047 | 27.5745 | <0.0001 |
| Egg temperature × Larval temperature | 1 | 461 | 2147225 | 166.063 | <0.0001 |
| Egg temperature × Sex | 1 | 461 | 678796 | 42.2873 | <0.0001 |
| Larval temperature × Sex | 1 | 461 | 85342.5 | 4.93496 | 0.026804 |
| Egg temperature × Larval temperature × Sex | 1 | 461 | 16796.2 | 1.07689 | 0.299938 |

**Figure 1:**
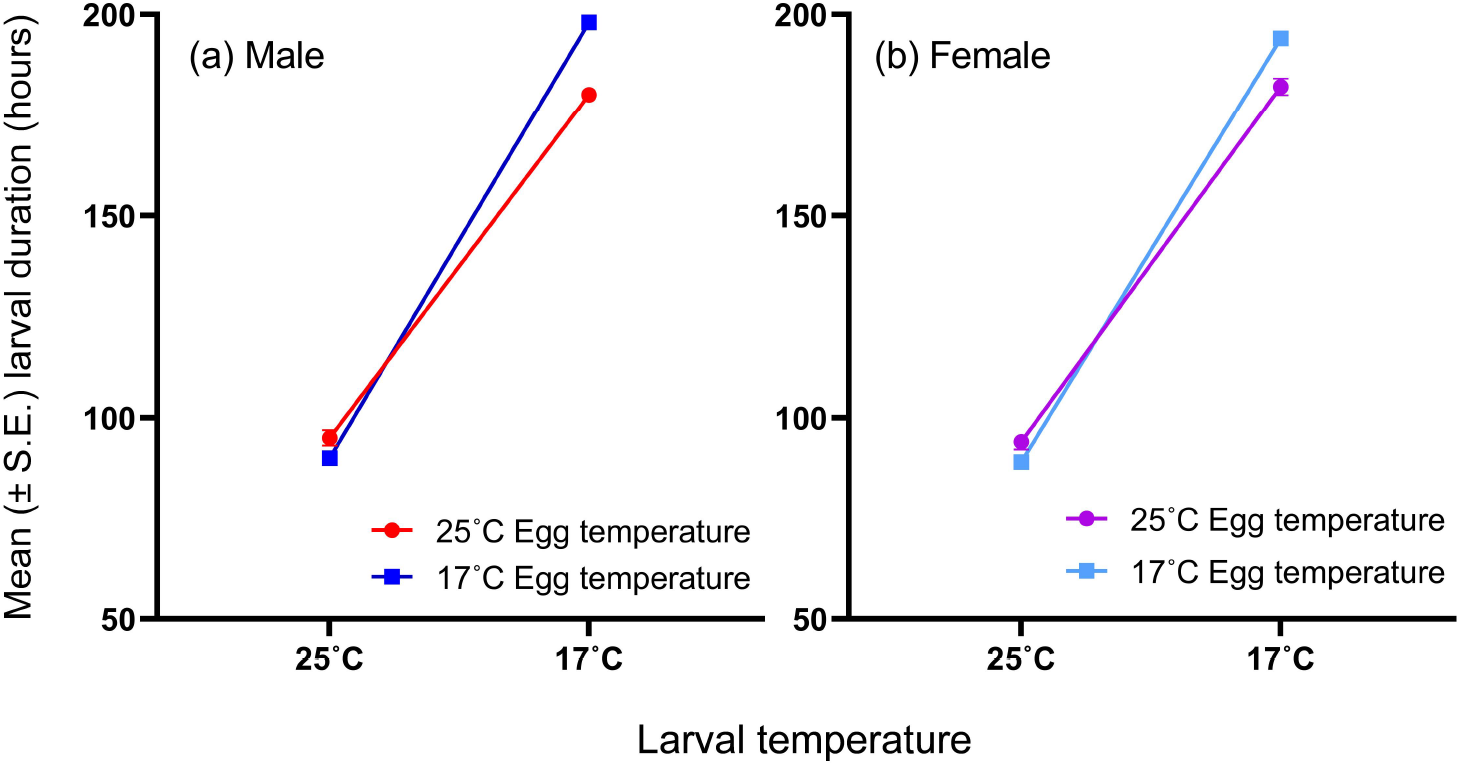
Reaction norm of mean larval duration across larval temperatures for a given egg stage temperature in (a) male and (b) female flies. The error bars in both plots represent standard error across replicate MCTs.

### Pupal duration

When pupae are reared at 25°C, their average development time ranges from 96 to 104 h across treatments. At 17°C mean pupal duration ranges from 172 to 201 h across treatments. Pupal duration shows significant effects of all three pre-adult stages and sex (*p* < 0.0001 for each) (Table 4). All two-way interactions involving egg larval and pupal temperatures are significant for pupal duration (*p* < 0.001 or 0.0001). However, the three-way interaction between egg larval and pupal temperatures reveals greater details (*p* < 0.0001) (Table 4).

**Table 4:** ANOVA table from aligned rank transform (ART) analysis showing nonparametric factorial effects of egg temperature, larval temperature, pupal temperature and sex on pupal duration.

| *Effect* | *df* | *df.res* | *SS* | *F* | *P* |
| --- | --- | --- | --- | --- | --- |
| Egg temperature | 1 | 453 | 2271449 | 172.548 | <0.0001 |
| Larval temperature | 1 | 453 | 2532872 | 200.069 | <0.0001 |
| Pupal temperature | 1 | 453 | 6058874 | 1312.78 | <0.0001 |
| Sex | 1 | 453 | 907893 | 57.9492 | <0.0001 |
| Egg temperature × Larval temperature | 1 | 453 | 604683 | 36.0436 | <0.0001 |
| Egg temperature × Pupal temperature | 1 | 453 | 2140315 | 159.691 | <0.0001 |
| Larval temperature × Pupal temperature | 1 | 453 | 299457 | 17.1511 | <0.0001 |
| Egg temperature × Sex | 1 | 453 | 191251 | 10.6788 | 0.001166 |
| Larval temperature × Sex | 1 | 453 | 820118 | 48.8803 | <0.0001 |
| Pupal temperature × Sex | 1 | 453 | 444640 | 26.7229 | <0.0001 |
| Egg temperature × Larval temperature × Pupal temperature | 1 | 453 | 382196 | 22.6091 | <0.0001 |
| Egg temperature × Larval temperature × Sex | 1 | 453 | 90393.9 | 5.01351 | 0.02563 |
| Egg temperature × Pupal temperature × Sex | 1 | 453 | 34886.9 | 1.9032 | 0.1684 |
| Larval temperature × Pupal temperature × Sex | 1 | 453 | 7956.81 | 0.49468 | 0.48221 |
| Egg temperature × Larval temperature × Pupal temperature × Sex | 1 | 453 | 90729.3 | 4.97527 | 0.026201 |

When pupae are developing at 25°C (Figure 2a), keeping the egg temperature constant (either 25°C or 17°C), but altering the larval temperature, significantly changes the pupal duration. At 25-17-25°C mean pupal duration changes to 100 h from 96 h at 25-25-25°C (*p* = 0.0002). At 17-17-25°C, average pupal duration is 104 h, which is significantly longer than average pupal duration at 17-25-17°C (98 h) (*p* = 0.0002). On the other hand, for pupae developing at 25°C, keeping the larval temperature constant (either 25 or 17°C), but changing the egg temperature also shows significant change in pupal duration. At 17-17-25°C mean pupal duration is 104 h while, at 25-17-25°C it is 100 h, and the difference is significant (*p* = 0.0009). The difference between mean pupal duration at 17-25-25°C and 25-25-25°C is only 2 h (98 *vs*. 96 h), and it as significant (*p* < 0.0001). ART ANOVA can detect small but consistent differences as significant, and we suspect this to be the case for the comparison between these two groups.” When pupae are developing at 17°C (Figure 2b), keeping the egg temperature constant (25°C), and altering the larval temperature decreases mean pupal duration of 201 h at 17-17-17°C to 195 h at 17-25-17°C. From 25-25-17°C to 25-17-17°C, the mean pupal duration increases from 172 h to 184 h. On the other hand, for 17°C pupal temperature, altering egg temperature while keeping larval temperature constant, changes the mean pupal duration of 201 h 17-17-17°C to 184 h at 25-17-17 h. Average pupal duration at 25-25-17°C is 172 h that extends to 195 h at 17-25-17°C. However, the pairwise comparisons did not detect any of these differences as significant. We suspect this may be due to slightly greater variability in development time at 17°C compared to 25°C, which could reduce the power of nonparametric tests to detect differences between groups.

**Figure 2a:**
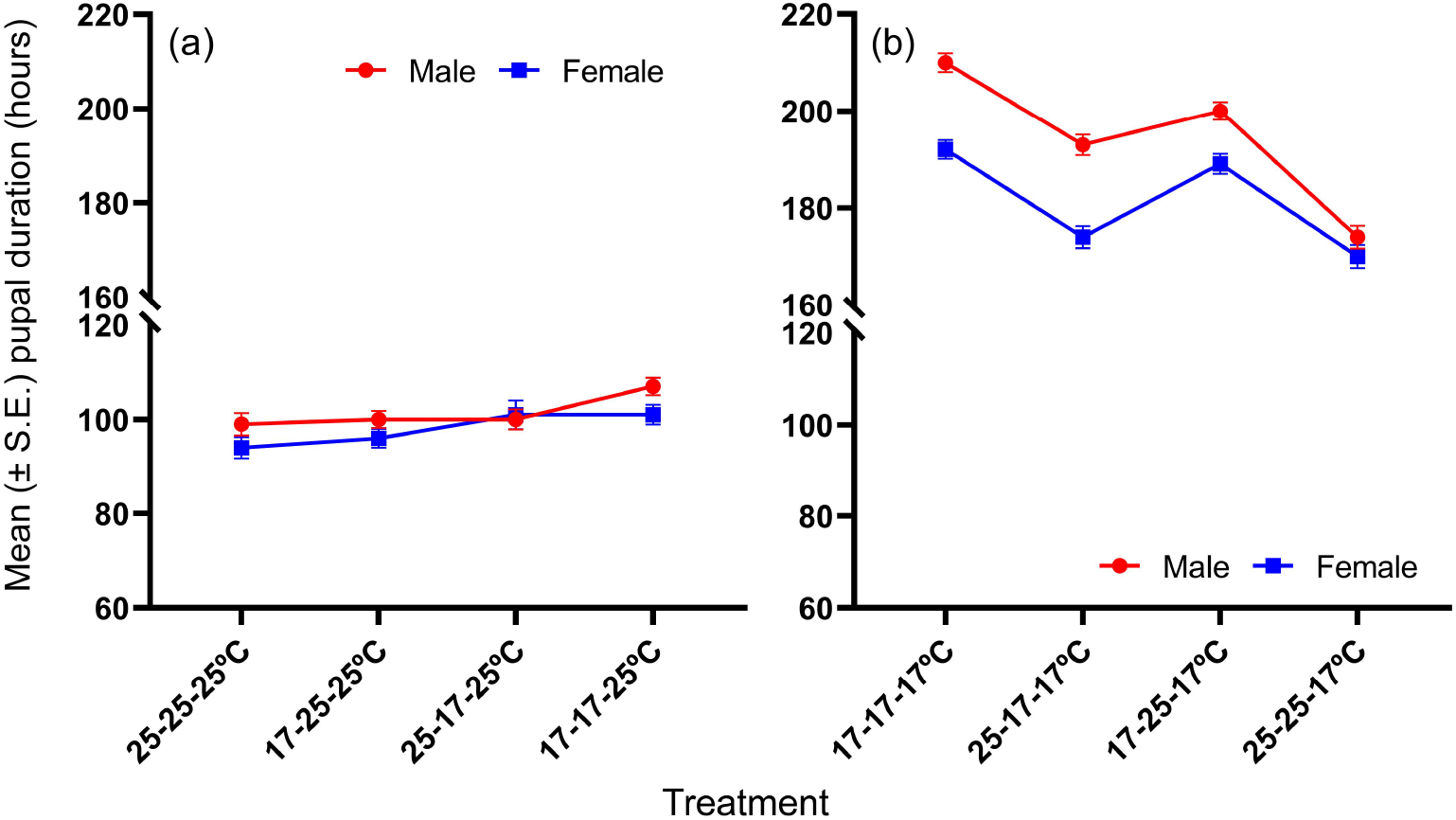
Mean pupal duration at 25°C across different thermal pre-treatments. The error bars represent standard error across replicate MCTs. Figure 2b: Mean pupal duration at 17°C across different thermal pre-treatments. The error bars represent standard error across replicate MCTs.

Average pupal duration in males (148 h) is significantly longer than that of females (142 h) (*p* < 0.0001). The interaction of pupal temperature and sex is significant because females show an 87% increase in average pupal duration from 25°C to 17°C while males show a 93% increase for the same (*p* < 0.0001). The four-way interaction of egg-larval-pupal temperatures and sex is significant (*p* = 0.0262), but post hoc comparisons show no significant difference between different groups.

### Egg to adult development time

Total or egg to adult development time is significantly influenced by all developmental stage temperatures and sex (*p* < 0.0001 for each) (Table 5). 17°C temperature during any stage significantly increases total developmental duration compared to 25°C.

**Table 5:** ANOVA table from aligned rank transform (ART) analysis showing nonparametric factorial effects of egg temperature, larval temperature, pupal temperature and sex on egg to adult development time.

| *Effect* | *df* | *df.res* | *SS* | *F* | *P* |
| --- | --- | --- | --- | --- | --- |
| Egg temperature | 1 | 453 | 5223961 | 758.663 | <0.0001 |
| Larval temperature | 1 | 453 | 6020071 | 1302.97 | <0.0001 |
| Pupal temperature | 1 | 453 | 6120664 | 1387.88 | <0.0001 |
| Sex | 1 | 453 | 1938412 | 140.633 | <0.0001 |
| Egg temperature × Larval temperature | 1 | 453 | 1380870 | 92.5074 | <0.0001 |
| Egg temperature × Pupal temperature | 1 | 453 | 1336785 | 92.2378 | <0.0001 |
| Larval temperature × Pupal temperature | 1 | 453 | 187510 | 11.3818 | 0.000805 |
| Egg temperature × Sex | 1 | 453 | 62754.1 | 3.56971 | 0.059481 |
| Larval temperature × Sex | 1 | 453 | 636798 | 40.3336 | <0.0001 |
| Pupal temperature × Sex | 1 | 453 | 846499 | 53.8916 | <0.0001 |
| Egg temperature × Larval temperature × Pupal temperature | 1 | 453 | 75627.6 | 4.53923 | 0.033665 |
| Egg temperature × Larval temperature × Sex | 1 | 453 | 37064.7 | 2.43229 | 0.119557 |
| Egg temperature × Pupal temperature × Sex | 1 | 453 | 7659.8 | 0.43298 | 0.510863 |
| Larval temperature × Pupal temperature × Sex | 1 | 453 | 119593 | 8.47546 | 0.003777 |
| Egg temperature × Larval temperature × Pupal temperature × Sex | 1 | 453 | 232648 | 13.5555 | 0.00026 |

All two-way interactions involving different developmental stage temperatures and three-way interaction of egg-larval-pupal temperatures are significant (*p* < 0.0001). Changing developmental temperature from constant 25°C to 17°C increases average egg to adult development time by 105% (*p* < 0.0001). Changing only egg temperature from 25°C to 17°C extends total development time by 6% (17-25-25°C *vs*. 25-25-25°C) (*p* < 0.0001) on average. Changing larval temperature from 25°C to 17°C increases total development time by 43% (25-17-25°C *vs*.25-25-25°C) (*p* < 0.0001) on average. Changing pupal temperature from 25°C to 17°C causes an average increase of 36% in total development time (25-25-17°C *vs*.25-25-25°C) (*p* = 0.004) (Figure 3a)

**Figure 3a:**
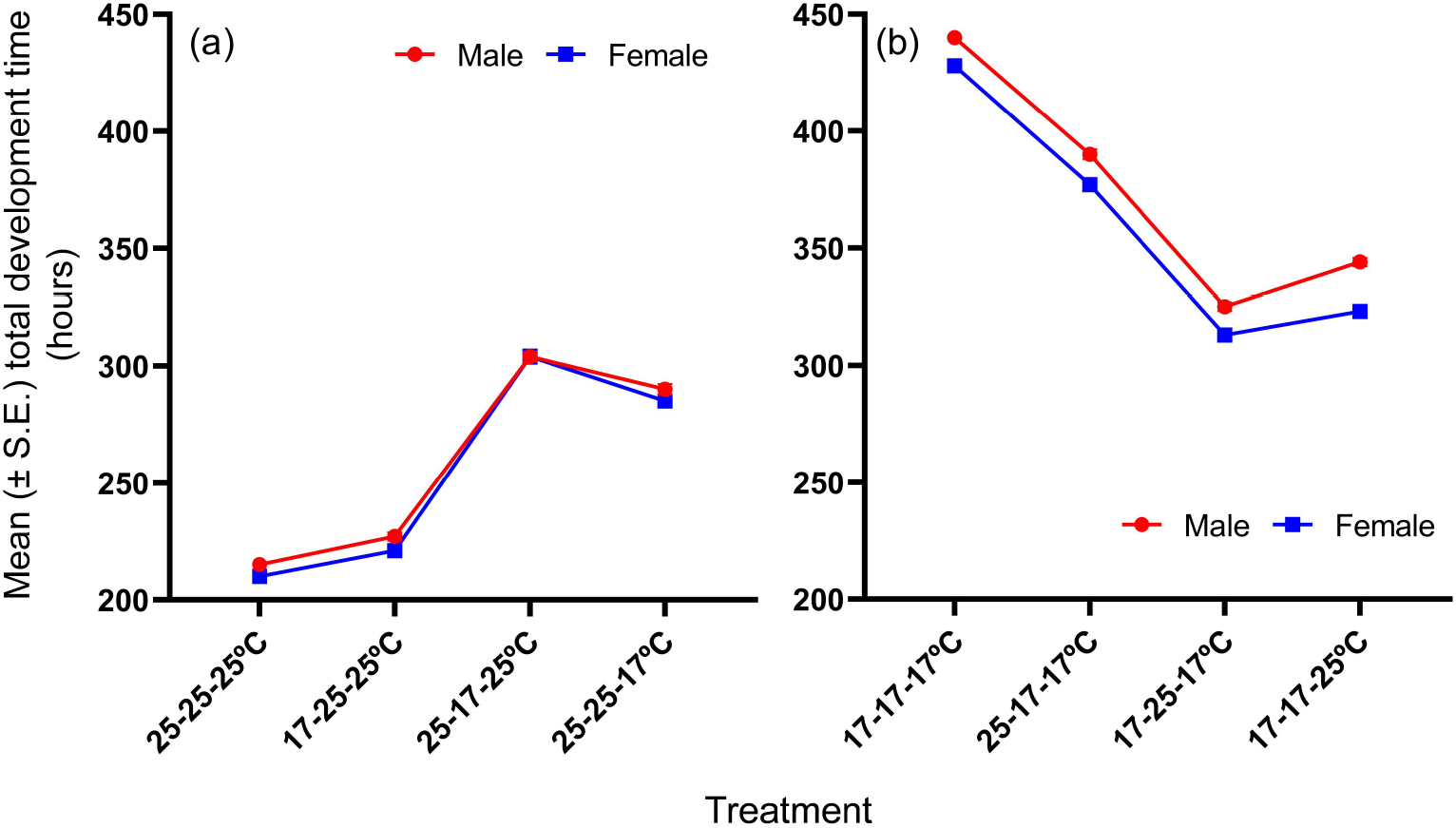
Mean sex-specific egg to adult development time across thermal treatments in which temperature is gradually changed from 25°C to 17°C across different life stages (first egg, then larva, then egg-larva both). The error bars represent standard error across replicate MCTs. Figure 3b: Mean sex-specific egg to adult development time across thermal treatments in which temperature is gradually changed from 17°C to 25°C across different life stages (first egg, then larva, then egg-larva both). The error bars represent standard error across replicate MCTs

Changing egg temperature from 17°C to 25°C decreases total development time by 12% (17-17-17°C *vs*. 25-17-17°C) on average. Changing larval temperature from 17°C to 25°C reduces total development time by 26% (17-17-17°C *vs*. 17-25-17°C) on average. The mean reduction in total development time Changing pupal temperature from 17°C to 25°C shortens total development time by 23% (17-17-17°C *vs*.17-17-25°C) (Figure 3b). However, none of these changes are significant. We suspect that because development time at each stage is longer at 17°C, variability is slightly higher than at 25°C, and as a result, post hoc comparisons on the transformed data do not detect significant differences among these groups.

Males have significantly longer total development time compared to females (*p* < 0.0001) (Figures 3a, 3b). Two-way interaction between egg temperature and sex is not significant, but those between larval temperature and sex, and pupal temperature and sex are significant (*p* < 0.0001 for each). Total development time of males is consistently longer than that of females across all groups. The interactions only reflect the slightly dissimilar degree of difference in thermal sensitivity of development time across sexes in some cases.

### Body size (wet weight)

Wet weight of the flies is significantly influenced by larval temperature, pupal temperature and sex (*p* < 0.0001 for each) (Figures 4a, 4b) (Table 6), but egg temperature does not have a main effect on the trait. 17°C rearing temperature during larval stage leads to higher wet weight at eclosion compared to 25°C (1.46 mg *vs*. 1.18 mg per fly). The interaction of egg and larval temperature is significant for wet weight (*p* < 0.0001) (Figures 5a, 5b). Pairwise comparisons reveal that the flies reared at 25°C egg temperature and 17°C larval temperature is significantly bigger than flies having 17°C at both stages (1.50 *vs*. 1.42 mg) (*p =* 0.01). Besides, the other shift temperature treatment, namely, 17°C egg temperature followed by 25°C larval temperature also results in formation of bigger flies than those grown at 25°C egg and larval temperatures (1.23 *vs*. 1.13 mg) (*p* < 0.0001). Flies grown at 17°C egg-larval temperature and 25°C egg-larval temperature are not significantly different (1.42 *vs*. 1.13 mg), and we suspect variability in data and rank transformation lead to lack of significant difference between the two groups. The difference between wet weight of flies having pupal stage temperature of 17°C *vs*. 25°C is small (1.33 mg *vs*. 1.30 mg), but ART ANOVA detects it as significant. We speculate this to be a result of transforming the data to ranks and ART detecting a minute but consistent difference as significant. The three-way interaction of egg-larval temperatures and sex is significant but post hoc comparisons show no significant difference between different groups.

**Table 6:**
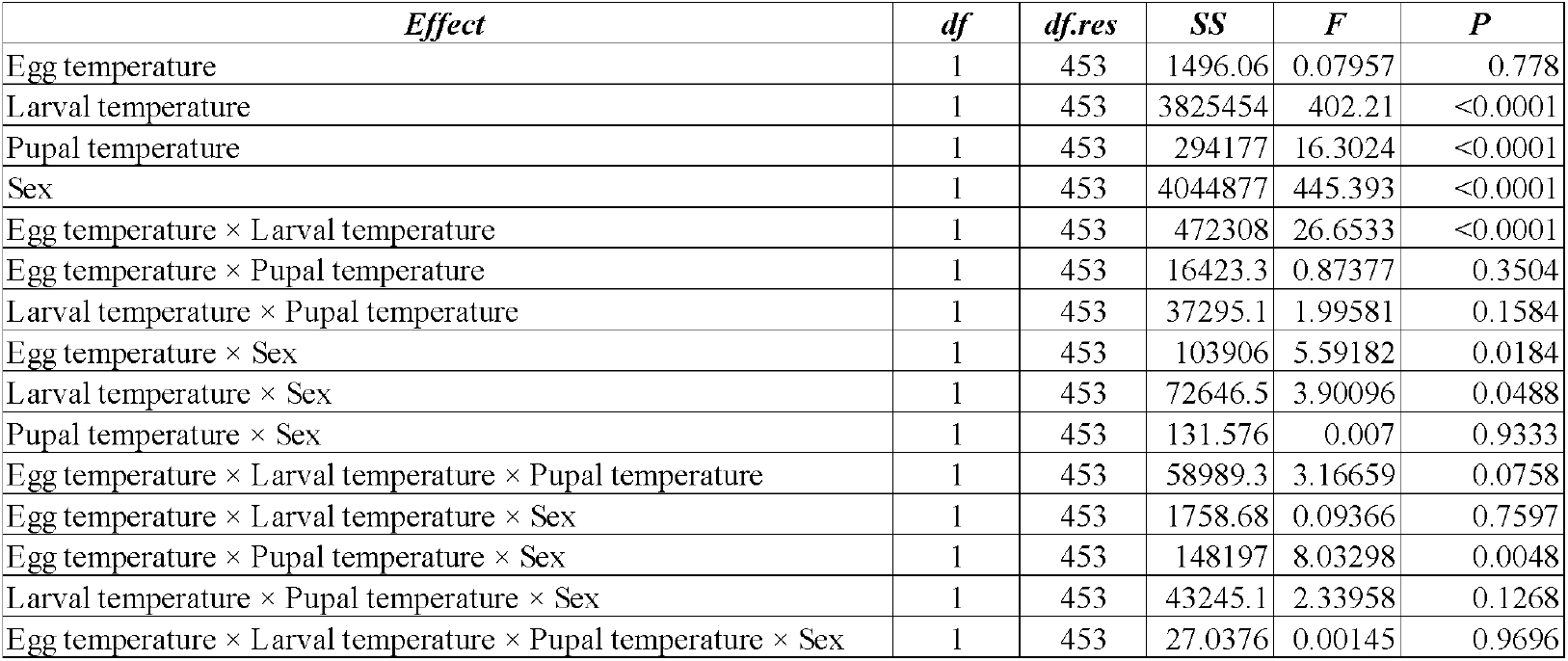
ANOVA table from aligned rank transform (ART) analysis showing nonparametric factorial effects of egg temperature, larval temperature, pupal temperature and sex on wet weight of flies.

| *Effect* | *df* | *df.res* | *SS* | *F* | *P* |
| --- | --- | --- | --- | --- | --- |
| Egg temperature | 1 | 453 | 1496.06 | 0.07957 | 0.778 |
| Larval temperature | 1 | 453 | 3825454 | 402.21 | <0.0001 |
| Pupal temperature | 1 | 453 | 294177 | 16.3024 | <0.0001 |
| Sex | 1 | 453 | 4044877 | 445.393 | <0.0001 |
| Egg temperature × Larval temperature | 1 | 453 | 472308 | 26.6533 | <0.0001 |
| Egg temperature × Pupal temperature | 1 | 453 | 16423.3 | 0.87377 | 0.3504 |
| Larval temperature × Pupal temperature | 1 | 453 | 37295.1 | 1.99581 | 0.1584 |
| Egg temperature × Sex | 1 | 453 | 103906 | 5.59182 | 0.0184 |
| Larval temperature × Sex | 1 | 453 | 72646.5 | 3.90096 | 0.0488 |
| Pupal temperature × Sex | 1 | 453 | 131.576 | 0.007 | 0.9333 |
| Egg temperature × Larval temperature × Pupal temperature | 1 | 453 | 58989.3 | 3.16659 | 0.0758 |
| Egg temperature × Larval temperature × Sex | 1 | 453 | 1758.68 | 0.09366 | 0.7597 |
| Egg temperature × Pupal temperature × Sex | 1 | 453 | 148197 | 8.03298 | 0.0048 |
| Larval temperature × Pupal temperature × Sex | 1 | 453 | 43245.1 | 2.33958 | 0.1268 |
| Egg temperature × Larval temperature × Pupal temperature × Sex | 1 | 453 | 27.0376 | 0.00145 | 0.9696 |

**Figure 4a:**
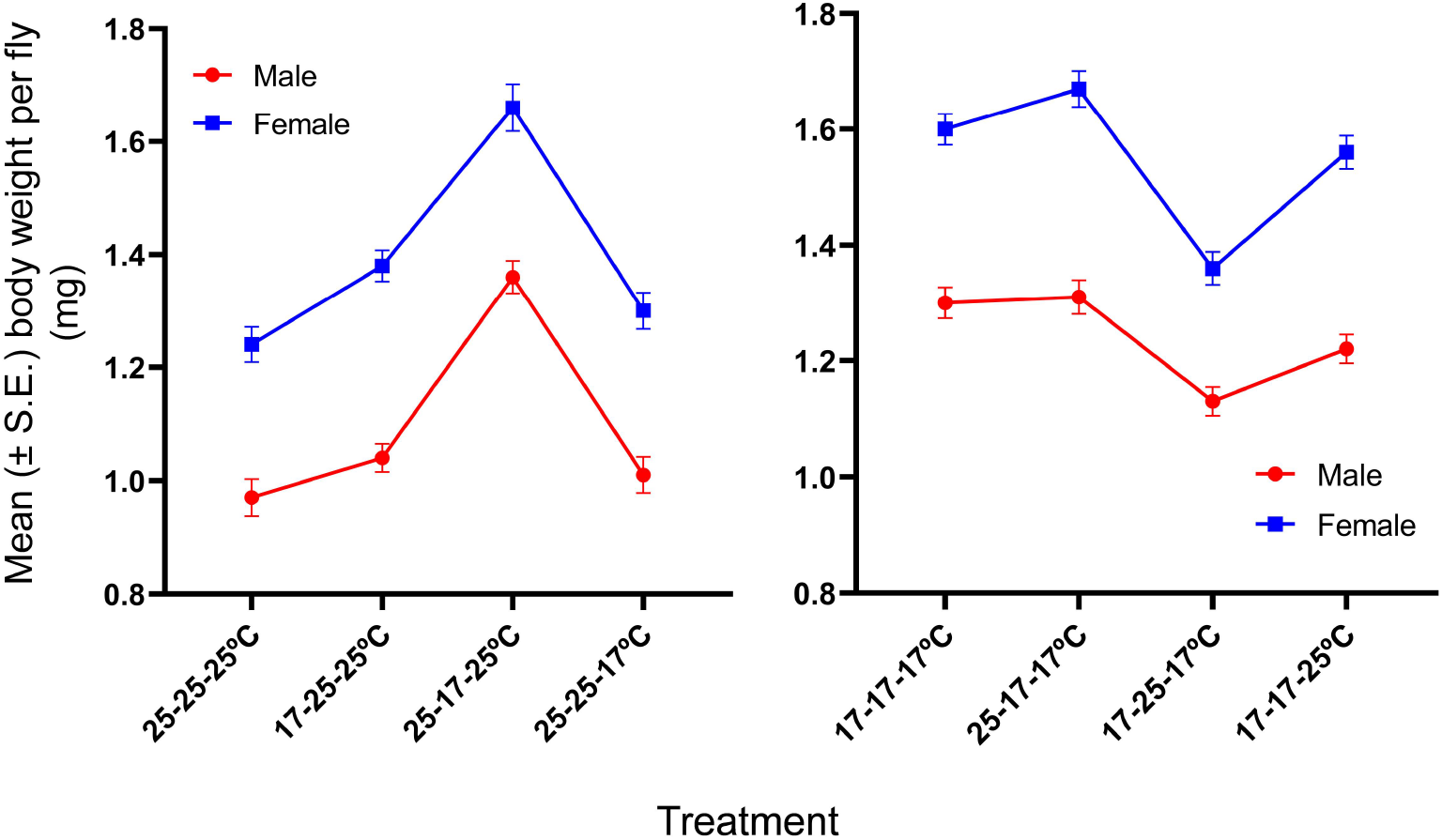
Mean sex-specific wet weight of flies across thermal treatments in which temperature is gradually changed from 25°C to 17°C across different life stages (first egg, then larva, then egg-larva both). The error bars represent standard error across replicate MCTs. Figure 4b: Mean sex-specific wet weight of flies across thermal treatments in which temperature is gradually changed from 17°C to 25°C across different life stages (first egg, then larva, then egg-larva both). The error bars represent standard error across replicate MCTs.

**Figure 5a:**
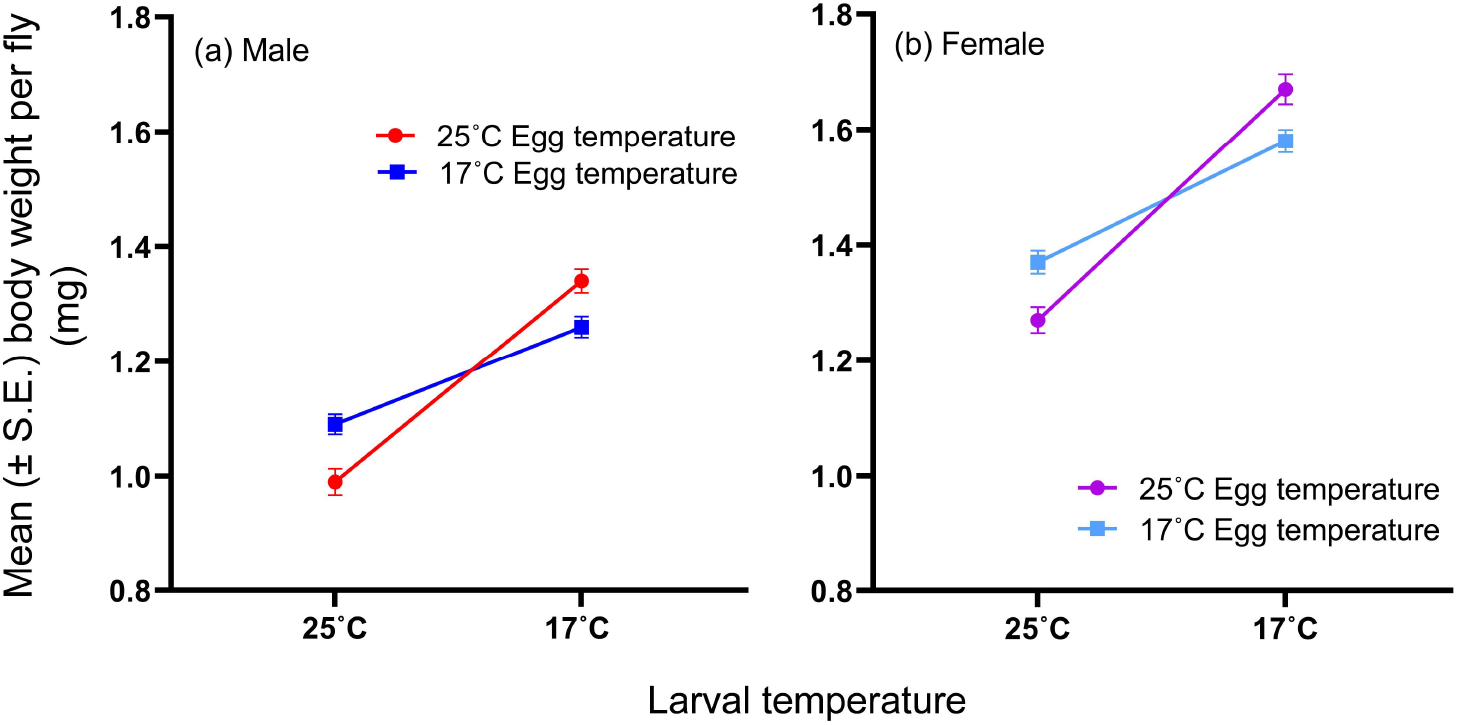
Mean wet weight of male flies for combination of egg and larval stage temperatures. The error bars represent standard error across replicate MCTs. Figure 5b: Mean wet weight of female flies for combination of egg and larval stage temperatures. The error bars represent standard error across replicate MCTs.

Females are significantly heavier than males (1.47 *vs*. 1.17 mg) (Figures 4a, 4b). Egg temperature × sex, larval temperature × sex, and egg temperature × pupal temperature × sex interactions are significant, but the pairwise comparisons do not show any significant difference. Other two- and three-way interactions and the four-way interaction are not significant.

## Discussion

### Effect of thermal shifts on development time

This study provides several important insights into how varying thermal conditions across the ontogeny influence developmental duration and final body size in flies. First, the larval and pupal durations show similar thermal plasticity as both get extended two-fold (100%) from 25°C to 17°C. The egg duration or larval hatching time increases by 70% in the same thermal range.

### Both low-to-high and high-to-low temperature shift at egg-larva transition accelerate subsequent larval development

When eggs are reared at 17°C instead of 25°C, average duration of the egg stage is lengthened by 15 hours (Supplementary figure S6). The difference in developmental speed set in by the egg stage temperature is carried onto the subsequent stages. When the egg-stage is at 25°C, but the hatched larvae are shifted to a cooler temperature of 17°C, the carryover effect of warmer egg temperature is observed during larval and development. Larvae at 17°C develop 8% faster on average compared to individuals developing at 17°C from the beginning (Figures 1a, 1b). This supports our *a priori* expectation that early exposure to a warmer temperature would leave an accelerating effect on subsequent development. (French et al., 1998; Gillooly et al., 2001; Zuo et al., 2011)

However, the reverse is not observed for larval development. Contrary to our prediction, exposure to a cold egg-stage temperature of 17°C does not prolong subsequent larval development. Instead, it results in a small but significant reduction in larval duration (Figures 1a, 1b). Thus, both upshifts and downshifts in temperature at the egg-to-larva transition leads to shorter larval development times. This aligns with some previous studies reporting modest acceleration of development under fluctuating thermal conditions (Roltsch et al., 1990, Al-Saffar et al., 1995,1996). Enzyme-catalyzed reaction rates are known to increase exponentially with temperature up to a thermal optimum (Somero, 1995; Hochachka and Somero, 2002). For organisms developing at suboptimal cold temperatures, enzymes likely operate with reduced catalytic efficiency. Studies suggest that when shifted to optimal temperatures, enzyme efficiency and metabolic flux in organisms may increase sharply (Fields et al., 2001; MacMillan et al., 2012; Chung and Schulte, 2020). We speculate this could be one of the reasons for the larval development becoming faster at 17°Cegg-25°Clarval rearing temperature compared to constant 25°C rearing temperature.

### Effect of egg and larval-stage temperatures on pupal duration

Carryover effects of earlier stage temperatures influence pupal development time in the expected directions.

For pupae developing at 25°C, altering only the larval stage temperature while keeping the egg temperature same, changes the pupal duration significantly reflecting clear carryover effect of larval stage temperature on pupal development time (Figure 2a). Although similar effect of thermal variation of larval temperature is observed for pupae developing at 17°C, and the magnitude of the mean change in pupal duration is much longer (Figure 2b), due to our experimental design and nonparametric tests we could not find a significant change in the latter case. Albeit the data clearly points towards the impact of larval temperature on pupal duration in both directions. Thus, a larval temperature of 17°C extends the subsequent pupal duration while 25°C larval temperature has an accelerating carryover effect on the trait (Figures 2a, 2b).

The effect of 17°C to 25°C egg-larval thermal shift is slightly more complex. This shift causes small but significant acceleration of larval development (as discussed before), while statistically it is found to increase the pupal duration. However, on closer look the mean increase in pupal duration from 25-25-25°C to 17-25-25°C is found to be only 1.3 h. Therefore, we infer while this shift leaves an accelerating effect on larval development, its impact practically does not extend up to the pupal stage (Figure 2a).

The carryover effect of a warm egg-stage temperature on the pupal duration is remarkable. Like larval temperature, the impact of egg temperature on pupal duration at 17°C could not be detected statistically, but it is observed in the data quite clearly as 25°C egg temperature has an overall reducing effect on the pupal duration compared to flies that were kept at 17°C as eggs. The accelerating effect of 25°C egg temperature on pupal duration at 25°C is clear and statistically significant as a thermal shift done for 25-17-17°C reared flies, both larval and pupal durations are significantly reduced compared to that of 17-17-17°C reared flies. This suggests a warm egg stage sets in a higher developmental rate spanning the rest of the ontogeny compared to a warm larval stage. While the thermal plasticity of any given developmental stage and its carryover effect on the subsequent stage was expected, we found the impact of egg stage temperature spanning the larval stage up to the pupal stage truly surprising. It suggests, for organisms who are exposed to fluctuating weather conditions during their development, developmental processes at a later stage may be contingent upon environmental conditions that occurred days ago and during a much earlier developmental stage (Angilletta et al., 2004; Kingsolver and Huey, 2008).

### Contribution of different pre-adult stages to thermal plasticity of total development time

We also found that variation in egg-stage temperature results in the smallest change in overall development time (Figures 3a, 3b), which is expected given that the egg stage is the shortest. In contrast, variability in larval or pupal temperatures has a more pronounced impact on total development time (Figures 3a, 3b). The larval and pupal stages are of comparable length and exhibit similar thermal plasticity when rearing temperature shifts from 25°C to 17°C. However, change in larval temperature has a greater effect on overall development time than change in pupal temperature (Figures 3a, 3b) This contrast arises because the larval stage occurs earlier in development, and thus any temperature induced changes to larval duration can exert a downstream influence on pupal development. While the carryover effect of larval temperature on pupal duration is small and not always statistically significant, the combined impact of altered larval and pupal durations driven by larval temperature shifts results in a substantial change in total development time.

### Development time in males vs. female flies

We found males have significantly longer pre-adult development time compared to females, and this is a well-established fact in *Drosophila. D. melanogaster* flies start mating within few hours of eclosion, and much of their sexual development occurs during the process of metamorphosis within pupae. (Casares et al., 1997; Whitworth et al., 2012; Testa et al., 2013) Literature suggests male flies take longer to attain sexual maturity and this causes longer duration of pupal development compared to females (Welbergen and Sokolowski, 1994; Nunney, 2007). However, our study indicates a small amount of delay in male development compared to females starts right from the larval stage. This is a novel finding as it suggests apart from sexual maturity there might be other sex-specific differences in development starting from larval stage that may have a small contribution in lengthening the development of male *Drosophila melanogaster*.

### Body size regulation under thermal shifts

#### The larval stage

Wet-weight data from our study indicate that, similar to constant-temperature conditions, the larval stage plays the most significant role in determining the thermal plasticity of adult body size under temperature-shift scenarios (Ghosh et al., 2013). This is expected because the larval period is relatively long and is the only feeding stage during fly development when most somatic growth occurs.

#### The pupal stage

The results indicate a main effect of pupal temperature on body size but the mean difference between the flies reared at contrasting temperatures is negligible. Therefore, we suspect this significance appears due to data transformation rather than reflecting a biologically relevant difference. Our previous work has also shown that the pupal stage does not contribute to thermal plasticity of size at eclosion (Ghosh et al., 2013). This is not surprising because the pupal stage is non-feeding and the fly undergoes metamorphosis during this stage within an enclosure, while losing some weight in the process (Ghosh et al., 2013). While the pupal development time is sensitive to temperature, the weight loss during pupal stage is similar at 17°C and 25°C (Ghosh et al., 2013).

#### The egg stage

There is limited information on how egg-stage temperature contributes to size regulation in flies. In a study by Hoover and Marks, 2021, fly cultures were subjected to single thermal shifts 24 hours after egg laying (from 18°C to 28°C and *vice versa*) and compared with flies reared at constant 18°C and 28°C. Interestingly, flies reared under the 28-18°C regime grew as large as those reared entirely at 18°C. Likewise, flies reared at 18-28°C also showed similar body size to the 18°C group. These results suggest that a brief early exposure to warm temperature does not impact final size if the majority of development occurs in cold. Conversely, early cold exposure followed by warm temperatures throughout the remainder of development significantly influenced adult size, highlighting a lasting effect of early cold condition. The study, therefore, indicates that the early egg-stage temperature may influence adult body size, but the effect may not be symmetric for warm *vs*. cold temperature. However, the experimental design of the study does not allow for precise evaluation of the specific role of egg-stage temperature on adult body size. In our study, as we investigated the effect of thermal upshifts and downshifts exactly at the egg-larva transition, we found that flies reared at 25°C during the egg stage followed by 17°C during the larval stage (25°Cegg-17°Clarval) are larger than those reared at 17°C (17°Cegg-17°Clarval) (Figures 4, 5a, 5b). The possible explanation for body size being bigger at 25-17°C is that a warmer egg-stage temperature presumably initiates a slightly higher early growth rate in the developing insect. This is consistent with our previous findings showing that larvae 24 hours post-hatching are larger at 25°C than at 17°C (Ghosh et al., 2013). Furthermore, our development time data demonstrate that a warm egg-stage temperature accelerates the rate of all subsequent stages. Taken together, our findings suggest that early exposure to warm temperatures may not only enhance *developmental rate* but also promote increased *growth rate* in *Drosophila*.

Regarding the effect of cold temperature during the egg stage, our *a priori* expectation was that exposure to a lower temperature during this period, followed by a warm larval phase, would reduce adult fly size compared to constant warm conditions. This was based on the assumption that early cold exposure would retard growth, which was supported by our earlier observaion that larvae from eggs reared at 17°C are smaller during early larval stages than those from eggs reared at 25°C (Ghosh et al., 2013). However, as mentioned before, our finding is the opposite. 17-25°C treatment not only results in reduced larval duration compared to constant rearing temperature of 25°C but also produces larger adult flies. This suggests that a thermal shift from a cooler egg stage to a warmer larval stage may lead to a modest increase in both developmental and growth rates, which possibly leads to a shorter larval duration and bigger flies compared to flies grown at warm egg and larval stages. Current models about thermal sensitivity of development and growth do not adequately explain such outcomes under changing temperature regimes (Czarnoleski et al., 2013; Manenti et al., 2021). A more detailed analysis of the thermal dynamics underlying differentiation, metabolism, and growth, integrating theoretical modes and empirical research, may help clarify these counterintuitive results.

Our findings indicate that the effect of egg-stage temperature on final body size is asymmetric when there is a temperature shift at the egg-larva transition. We speculate that a warm egg stage may elevate growth rates slightly, not only during the egg stage but also during the subsequent cold larval stage, compared to constant cold rearing-suggesting that the growth advantage begins *prior* to the thermal shift. Conversely, when the egg stage is cold, the shift to a warmer larval environment may trigger changes in developmental and growth dynamics, resulting in flies that are ultimately larger than those reared under constant warm conditions. In this scenario, the size difference between constant warm and cold egg-warm larval rearing likely emerges *after* the temperature shift. Taken together, our results suggest that egg-stage temperature can influence not only early larval size but also regulate overall growth trajectories in *Drosophila*, particularly under fluctuating thermal conditions.

## Conclusion

The unique experimental design of this study allowed us to (a) investigate how the thermal environment of one pre-adult stage interacts with that of subsequent stages to shape developmental durations, and (b) precisely quantify the contribution of each life stage to the body size plasticity in *Drosophila melanogaster* under variable thermal environments. Together, these findings highlight that under variable temperatures, thermal responses emerge from a network of stage-specific sensitivities, cross-stage carryover effects, and nonlinear interactions between development and growth.

We show that development during any pre-adult stage is shaped not only by the ambient temperature but also by the thermal conditions experienced in the the past including thermal environment experienced several days earlier during an early developmental period like the egg stage can exert its effect on duration of developmental stages that occur much later, like the pupal stage. We found that the larval and pupal durations in flies are of comparable length and exhibit high thermal plasticity, while the relatively shorter egg stage has slightly lower plasticity. Although the larval and pupal stages have similar durations and comparable thermal plasticity, the larval temperature exerts a stronger influence on total developmental time. This is likely because thermal conditions during larval development not only determine larval duration but also leave a carryover effect that extends into and alters pupal duration.

Another important finding from the study is contrary to the simple expectation that a warmer temperature during an earlier stage would accelerate subsequent stages while a colder temperature would cause only a slowdown, our results demonstrate that the interactions between developmental stages and their thermal environments are far more complex. We show that a higher egg stage temperature may reduce developmental duration in subsequent cooler environments, but can paradoxically result in slightly larger flies due to enhanced early growth rates. Interestingly, a cold egg stage followed by a warmer larval stage can also shorten development time yet increase final body size compared to constant warm rearing. The exact mechanism for this phenomenon remains to be explored.

Additionally, while our study supports the established view that differences in development time among male and female flies arise primarily from differences in pupal duration, our results also indicate that males experience a slight developmental delay beginning as early as the larval stage. Another important finding of this study is that although the larval stage remains the primary determinant of adult body size, the egg-stage temperature also contributes to some extent. The contribution of pupal stage temperature on thermal plasticity of body size is negligible.

Several important directions emerge from this work. First, identifying the physiological and molecular mechanisms that mediate cross-stage thermal carryover effects remains a crucial next step. For example, investigating whether thermal shifts alter endocrine signaling, metabolic rates, energy utilization, or nutrient assimilation during development could illuminate why certain combinations of early and late temperatures accelerate development but still allow larger body sizes. Second, future experiments that manipulate temperatures dynamically, such as brief pulses, ramped thermal changes, or ecologically realistic fluctuating regimes, would help determine whether the patterns observed here generalize to more naturalistic conditions. Third, our findings raise intriguing questions about whether similar cross-stage interactions occur in other ectotherms. Comparative work across species with different life histories could clarify whether these carryover effects represent a general feature of ectothermic development or reflect *D. melanogaster* specific physiological responses. Finally, understanding how thermal history shapes body size and developmental timing has implications for predicting organismal responses to climate change. As natural thermal environments become increasingly variable, the ability of early thermal experiences to modulate later developmental outcomes may strongly influence population dynamics, fitness, and thermal adaptation.

## Author contributions

AC: design of the study, acquisition of data, analysis of data, validation, drafting the article, revising the article critically for important intellectual content; RR: design of the study, acquisition of data, analysis of data, validation, revising the article critically for important intellectual content; PB: acquisition of data SG: conception and design of the study, analysis and interpretation of data, supervision, drafting the article, revising the article critically for important intellectual content, resources, funding acquisition. All authors approved the version submitted.

## Acknowledgements

We thank Sreebes Deb Sharma, Abhisha Dutta, Sohang Pal, Jayalakshmi D. and Adarsh Tripathi for fly population maintenance and media preparation, Ranjit Pradhan and Manas Ranjan Mallik for general help in the laboratory.

## Data availability

The raw data used in the study, and custom R scripts written and used for data analyses are available in the supplementary material.

## Declaration of generative AI in scientific writing

During the preparation of this work generative AI has not been used.

