## Supplementary file for "Thermal Plasticity of Stage-specific Development Time and Adult Body Size under Temperature Shifts: A Case Study Using *Drosophila melanogaster*"

(supplementary material)

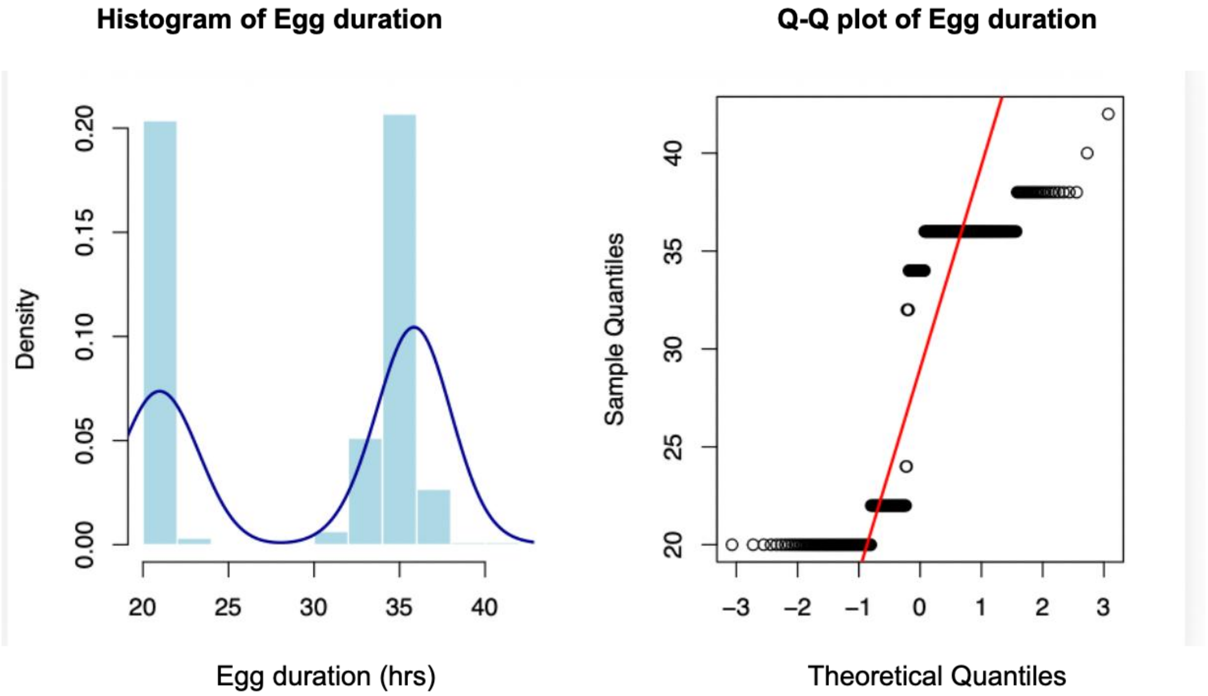


Figure S1: Density histogram and Q-Q plot of normality assessment for egg duration

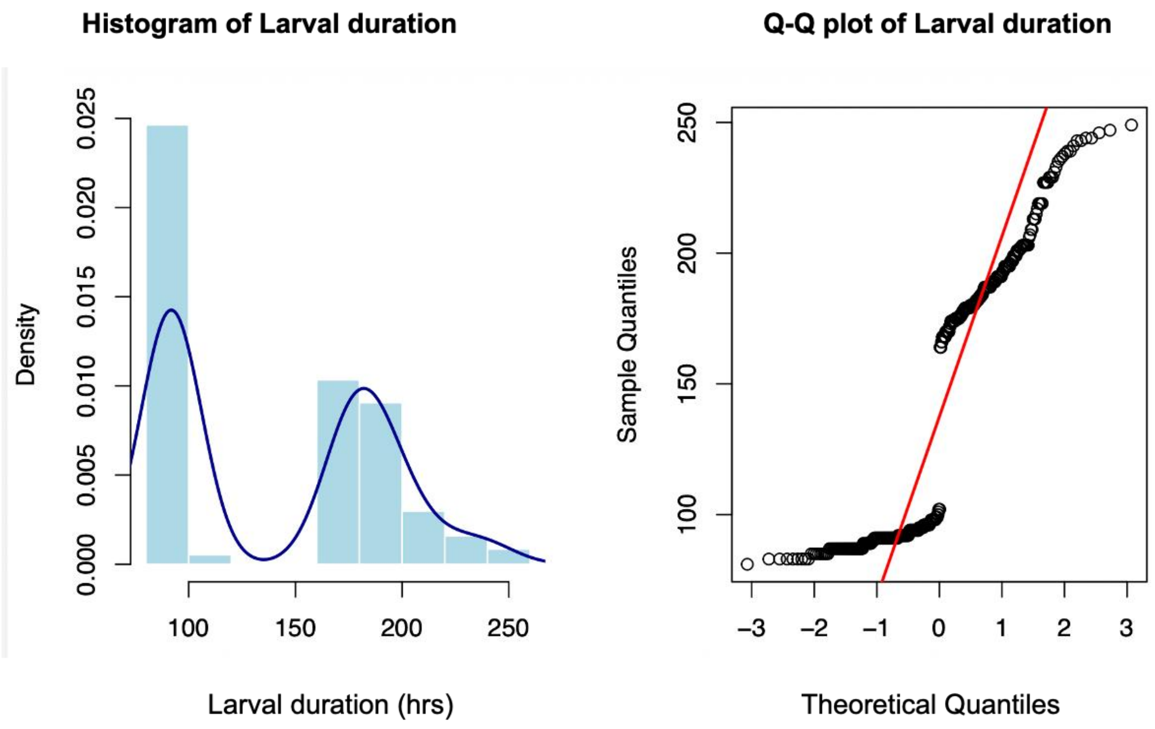


Figure S2: Density histogram and Q-Q plot of normality assessment for larval duration

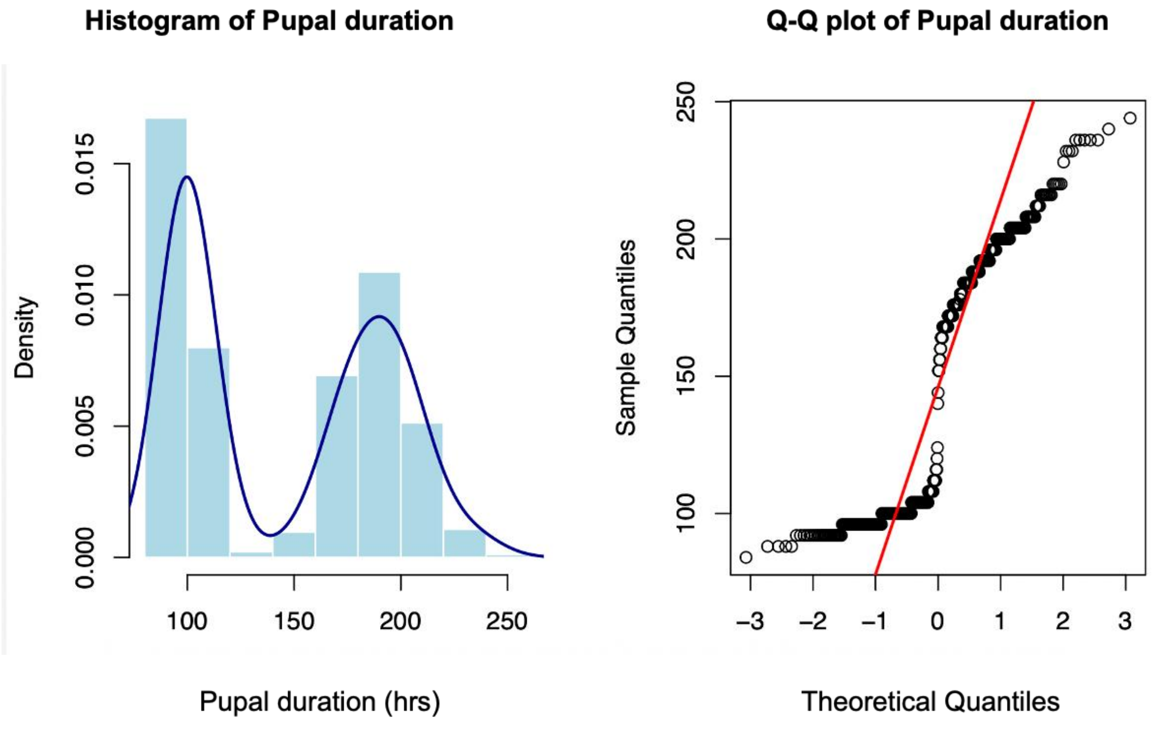


Figure S3: Density histogram and Q-Q plot of normality assessment for pupal duration

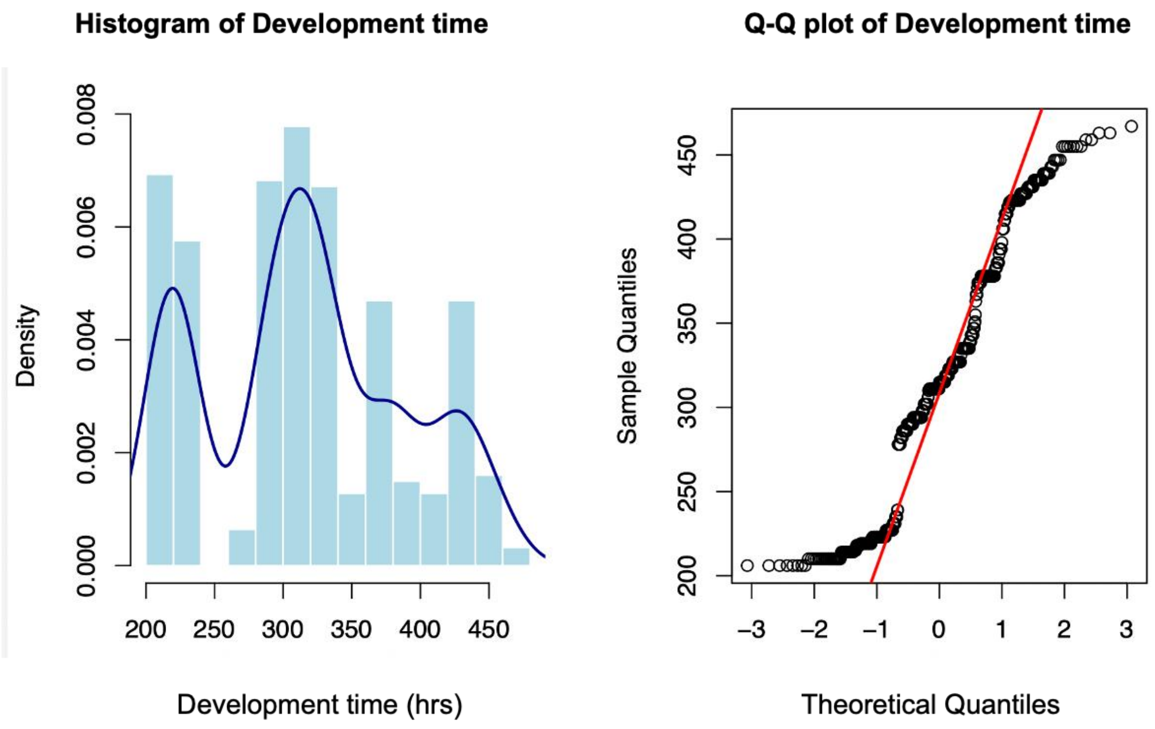


Figure S4: Density histogram and Q-Q plot of normality assessment for total development time

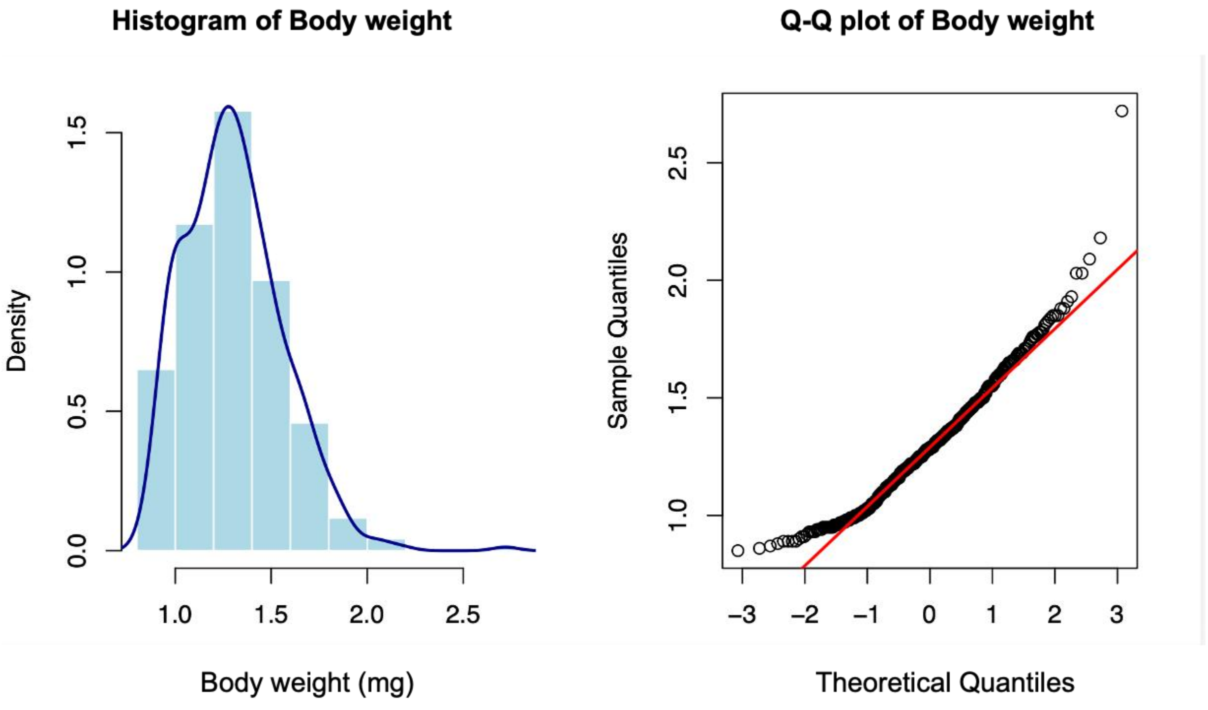


Figure S5: Density histogram and Q-Q plot of normality assessment for Body weight of the flies

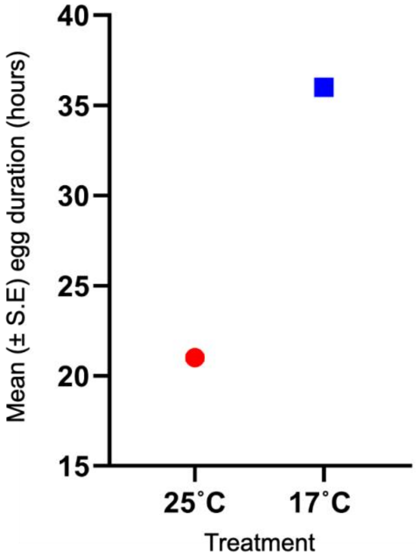


Figure S6: Mean (± S.E.) egg duration at 25°C and 17°C

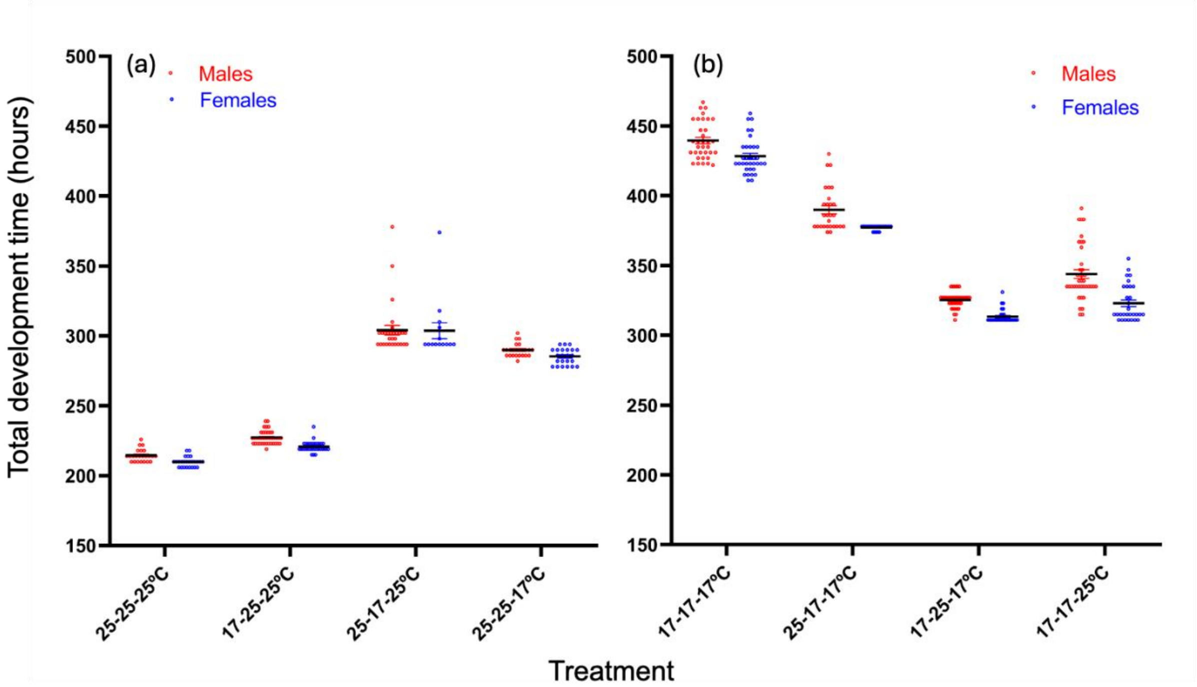


Figure S7: Sex specific egg to adult development time of individual flies across all thermal treatments.

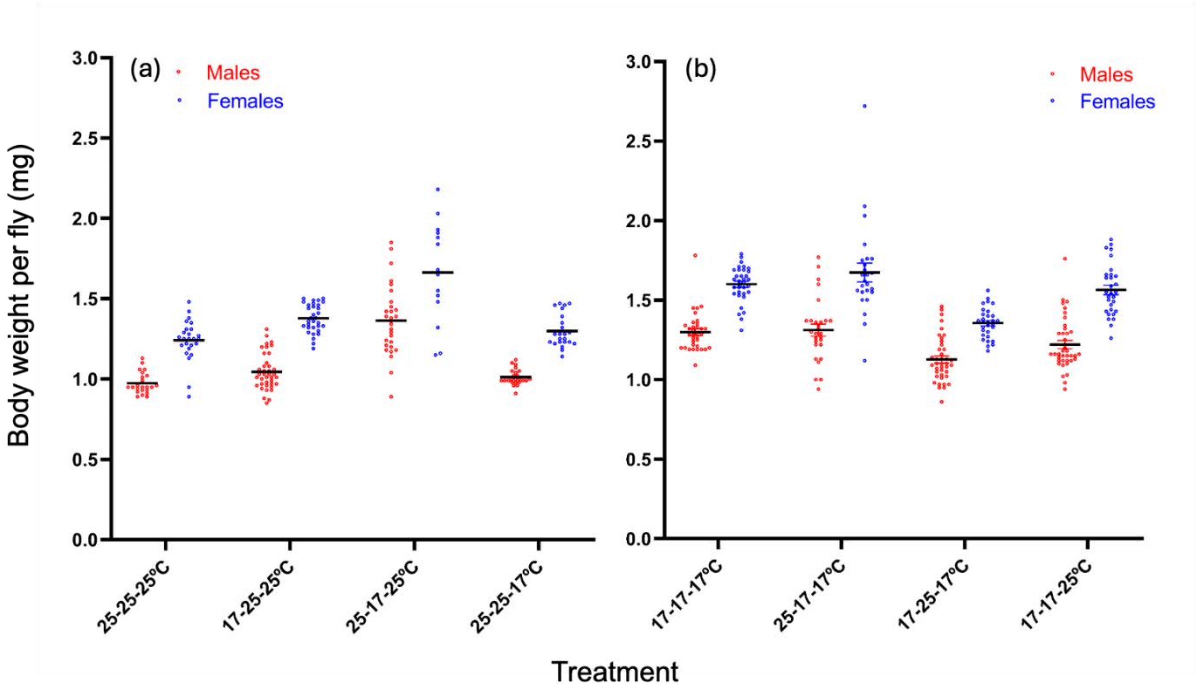


Figure S8: Sex specific wet weight of individual flies across all thermal treatments.
